# Membrane Fusion can be Driven by Sec18/NSF, Sec17/αSNAP, and *trans*-SNARE complex without HOPS

**DOI:** 10.1101/2021.08.06.455435

**Authors:** Hongki Song, William Wickner

## Abstract

Yeast vacuolar membrane fusion has been reconstituted with R, Qa, Qb, and Qc-family SNAREs, Sec17/αSNAP, Sec18/NSF, and the hexameric HOPS complex. HOPS tethers membranes and catalyzes SNARE assembly into RQaQbQc *trans*-complexes which zipper through their SNARE domains to promote fusion. Previously, we demonstrated that Sec17 and Sec18 can bypass the requirement of complete zippering for fusion (Song et al., 2021), but it has been unclear whether this activity of Sec17 and Sec18 is directly coupled to HOPS. HOPS can be replaced for fusion by a synthetic tether when the three Q-SNAREs are pre-assembled. We now report that SNARE zippering-arrested fusion intermediates that are formed without HOPS support Sec17/Sec18-triggered fusion. This zippering-bypass fusion is thus a direct result of Sec17 and Sec18 interactions: with each other, with the platform of partially zippered SNAREs, and with the apposed tethered membranes. As these fusion elements are shared among all exocytic and endocytic traffic, Sec17 and Sec18 may have a general role in directly promoting fusion.

---

Intracellular membrane fusion is catalyzed by protein families which are conserved from yeast to humans and among the organelles (Wickner and Rizo, 2017). These include Rab-family GTPases, large tethering complexes which bind to Rabs (Baker and Hughson, 2016), membrane-anchored SNARE proteins which assemble into *trans-*complexes that bridge membranes before fusion, and SNARE chaperones of the SM, Sec17/αSNAP and Sec18/NSF families. Yeast vacuole fusion combines Rab-effector tethering and SM functions into the large hexameric HOPS (**<u>ho</u>**motypic fusion and vacuole **<u>p</u>**rotein **<u>s</u>**orting) complex (Wurmser et al., 2000; Seals et al., 2000; Stroupe et al., 2006). HOPS binds to acidic lipids (Orr et al., 2015) and to its Rab Ypt7 on each fusion partner membrane (Hickey et al., 2010), activating it (Torng and Wickner, 2020) to initiate N-to C-directional assembly among the R, Qa, Qb, and Qc SNAREs. Though Sec17 and Sec18 will block fusion from spontaneously assembled *trans*-SNARE complexes, HOPS confers resistance to Sec17 interference (Mima et al., 2008). Without Sec17 or Sec18, HOPS-assembled *trans*-SNARE complexes require complete zippering to induce fusion (Schwartz and Merz, 2009). The ubiquitous chaperones Sec17 and Sec18 will bind to partially zippered SNARE complexes and use the N-terminal apolar loop on Sec17 to trigger fusion without needing complete zippering (Schwartz and Merz, 2009; Zick et al., 2015; Schwartz et al., 2017; Song et al., 2021). Sec17 has direct affinity for HOPS (Song et al., 2021); it has been unclear whether HOPS is needed for Sec17 and Sec18 to engage partially zippered SNAREs and mediate zippering-bypass fusion.

We now exploit a synthetic tether to show that HOPS is not required for Sec17 and Sec18 to drive zippering-bypass fusion. The 4 SNAREs, Sec17/αSNAP, and Sec18/NSF, which are the fundamental components of the 20s particle (Zhao et al., 2015), suffice to drive fusion. As a synthetic tether, we employ the dimeric protein glutathione-S-transferase (GST) fused to a PX domain. Dimeric GST-PX binds PI3P to tether membranes bearing PI3P (Song and Wickner, 2019). When Q-SNAREs are pre-assembled into a QaQbQc ternary complex on one fusion partner membrane, tethering by GST-PX supports fusion without the need for SM function (Song and Wickner, 2019). This fusion relies on SNARE zippering, as it is blocked by deletion of the C-terminal region of the Qc SNARE domain, the Qc3Δ mutation (Schwartz and Merz, 2009; Song and Wickner, 2019). We find that GST-PX tethered membranes bearing R and QaQbQc3Δ SNAREs on fusion partners, unable to completely zipper and fuse because of the shortened Qc SNARE domain, are rescued from this arrested state and will fuse upon addition of Sec17 and Sec18. Thus the Sec17 and Sec18 fusion functions do not rely on interactions with HOPS, instead acting through their interactions with each other, with tethered membranes, and with a partially-zippered *trans*-SNARE binding platform.

## Results

Proteoliposomes were prepared with vacuolar lipids, with membrane anchored Ypt7, and with either the R- or the 3Q-SNAREs. The Qc SNARE was either wild-type with its full-length SNARE domain or Qc3Δ which lacks the C-terminal 4 heptads of its SNARE domain. The Ypt7/R- and Ypt7/3Q-proteoliposomes bore lumenal fusion-reporter fluorescent proteins, either Cy5-labeled streptavidin or biotinylated phycoerythrin (Figure 1A). Proteoliposomes were purifed by flotation to remove unincorporated proteins. When these proteoliposomes are mixed, their lumenal fluorescent proteins are separated by at least the thickness of two lipid bilayers, too far for measurable fluorescence resonance energy transfer (FRET). Upon fusion and the attendant content mixing, the binding of biotin to streptavidin brings the Cy5 and phycoerythrin fluorophores into intimate contact, yielding a strong FRET signal (Zucchi and Zick, 2011). Fusion incubations were performed with mixed Ypt7/R and Ypt7/3Q proteoliposomes and with external nonfluorescent streptavidin to block any signal from proteoliposome lysis. Each incubation had either HOPS or GST-PX to tether the membranes (Figure 1A). Also present from the start were either a) buffer alone, b) Sec17, c) Sec18 and ATPγS, or d) both Sec17 and Sec18/ATPγS. Fusion was monitored by FRET between the lumenal probes; the initial rate during the first 5 minutes is termed the α portion in Figures 1B-E. At 30 minutes, each reaction received a supplement of the components not added at time 0, i.e. a) Sec17, Sec18 and ATPγS, b) Sec18 and ATPγS, c) Sec17, or d) buffer alone, so that all incubations had Sec17, Sec18, and ATPγS as the incubation continued in the β portion of the experiment (Fig1Bβ-Eβ), from 30 minutes to 32 minutes. Distinct information can be gleaned from the α and β intervals of the experiment, and these are considered in turn below.

**Figure 1.**
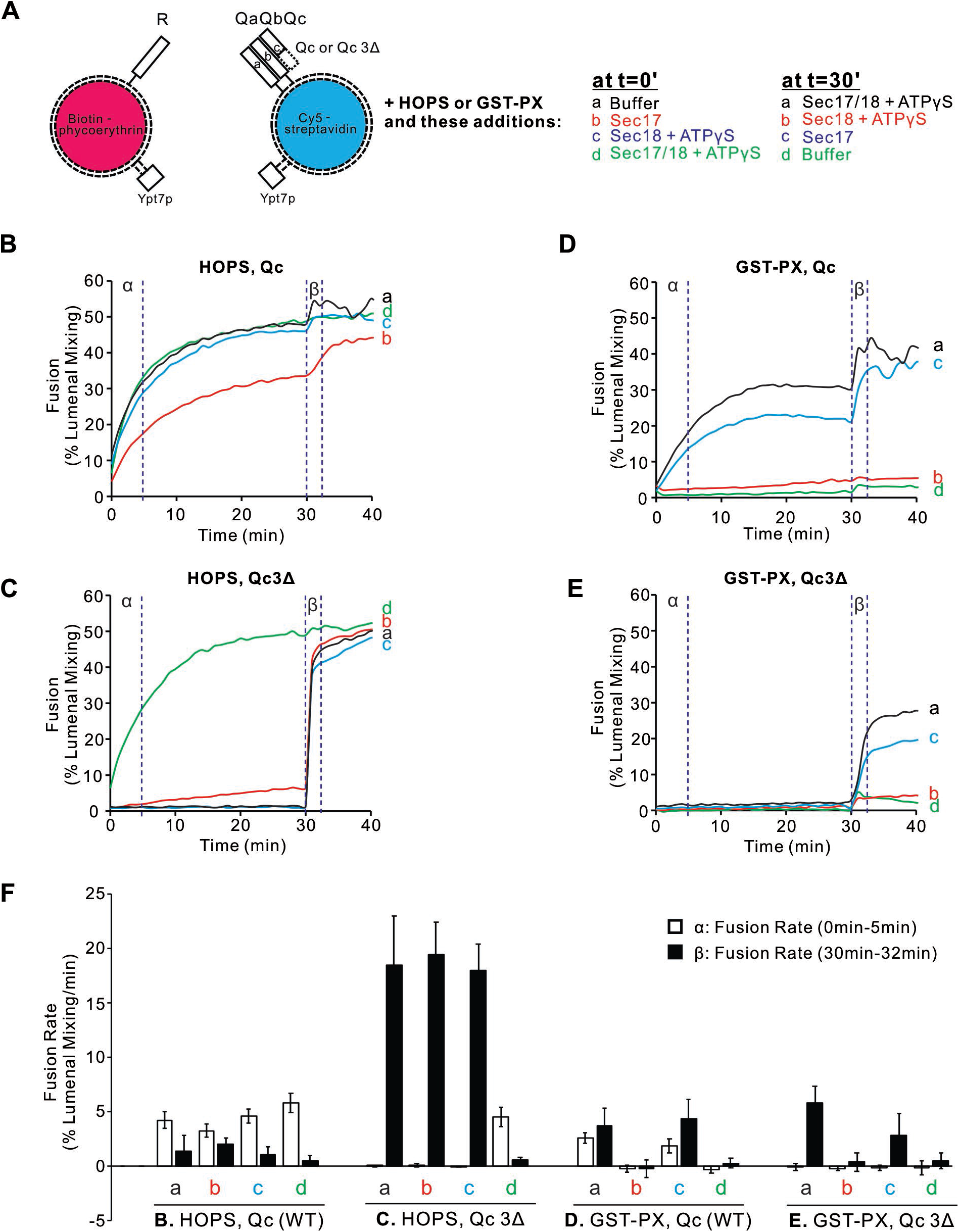
Sec17, Sec18, and ATPγS promote rapid and HOPS-independent fusion without SNARE zippering. Proteoliposomes were prepared with vacuolar lipids, membrane-anchored Ypt7, and either R or the 3 Q-SNAREs with molar ratios of 1Ypt7:8,000 lipids and 1 of each SNARE/16,000 lipids. Since Qc3Δ is a labile member of SNARE complexes (Song et al., 2021), we used a 5-fold molar excess to the other SNAREs in preparing Ypt7/QaQbQc3Δ proteoliposomes. The initial mixtures at t=0 were 18 μl, and remaining components were added at 30min in 2μl. A vertical bar separates the first 30min, termed α, and the 2-minute β interval after further additions. Experiments were repeated in quintuplicate; mean values and standard deviations are shown.

### HOPS is not required for Sec17/Sec18 stimulation of fusion

When the 3Q complex includes wild-type full-length Qc, HOPS-mediated fusion (Figure 1Bα, black curve) shows only minor effects from adding either Sec17 (red), Sec18 with a nonhydrolyzable ATP analog (blue), or Sec17, Sec18, and ATPγS (green). In contrast, when the 3Q complex includes Qc3Δ to arrest SNARE zippering and block fusion (Figure 1Cα, black curve), both Sec17 and Sec18 are required to bypass the zippering-arrest and allow fusion (Figure 1Cα, contrast the green curve vs the blue, red or black curves).

The dimeric tether GST-PX (Song and Wickner, 2019) also supports fusion with pre-assembled wild-type Q-SNAREs (Figure 1Dα, black curve), but this fusion is blocked by Sec17 (Figure 1Dα, red curve) without rescue by Sec18 (green curve). When fusion with the dimeric GST-PX tether is blocked by the Qc3Δ mutation (Figure 1Eα, black curve), there is no rescue by Sec17, alone or in combination with Sec18/ATPγS (Figure 1Eα, red and green curves), since HOPS is the only tether which bypasses inhibition by Sec17 (Song and Wickner, 2019).

At the end of these 30 minute incubations, each reaction received a further addition of any components not added at time 0. After this addition, each incubation had Sec17, Sec18, and ATPγS. Fusion incubations continued in the β portion of the experiment, from 30 minutes to 32 minutes, and beyond. Although full-length SNAREs support zippering and fusion with either the

HOPS or GST-PX tether, a kinetic intermediate accumulates which gives some additional fusion upon addition of Sec17/Sec18/ATPγS (Figure 1, Bβ and Dβ, black curves). When SNARE zippering and the attendant fusion was blocked by the Qc3Δ mutation, HOPS-dependent zippering-bypass fusion requires Sec17, Sec18, and ATPγS (Figure 1Cα), as reported (Song et al., 2020; 2021). In their absence, fusion intermediate accumulated, since there is rapid fusion upon their addition (Figure 1Cβ, curves a-c). Strikingly, though the presence of Sec17 from the start of the incubation blocks the formation of rapid-fusion intermediate with the GST-PX tether (Figure 1Eβ, red curve), rapid-fusion intermediate does accumulates with the GST-PX tether when Sec17 is absent, as Sec17 addition at 30minutes triggers fusion (Figure 1Eβ, curves a and c). Replicates of this experiment were quantified for the fusion rate during the α and β intervals (Figure 1F). Inhibition by Sec17 with the GST-PX tether instead of HOPS is only seen when Sec17 is present from the start of the incubation, prior to *trans*-SNARE assembly (Figure 1E, red curve b and green curve d), but once membranes have undergone tethering and *trans*-SNARE assembly, Sec17 and Sec18 support zippering-bypass fusion without HOPS (Figure 1Eβ, blue and black curves).

## Discussion

Sec17 has direct affinity for SNAREs and can inhibit fusion of the organelle (Wang et al., 2000) or block fusion of reconstituted proteoliposomes without HOPS (Mima et al., 2008). HOPS engages each of the 4 SNAREs (Stroupe et al., 2006; Baker et al., 2015; Song et al., 2020) and catalyzes their assembly (Baker et a., 2015; Orr et al., 2017; Jiao et al., 2018; Song et al., 2020) in a manner which renders fusion relatively resistant to Sec17 (Mima et al., 2008; Song and Wickner, 2019). Excessive Sec17 blocks the initial stages of fusion while stimulating fusion when added after *trans*-SNARE assembly (Zick et al., 2015). With the synthetic tether GST-PX, Sec17 which interacts with SNAREs before they assemble into *trans*-complexes inhibits the subsequent fusion (Song and Wickner, 2019). We now show that once fusion intermediates of *trans*-SNARE complexes have formed, Sec17 and Sec18 will promote efficient zippering-bypass fusion whether tethering is by HOPS or by the synthetic tether GST-PX (Figure 1Cβ and Eβ).

HOPS serves both as a tether (Hickey et al., 2010) and catalyst to initiate SNARE assembly (Baker et al., 2015), but it has been unclear whether it catalyzes the later stages of zippering or is needed for Sec17/Sec18-induced fusion without zippering. The engagement of SNAREs by HOPS (Stroupe et al., 2006; Baker et al., 2015; Song et al., 2020) largely bypasses Sec17 inhibition. Once SNAREs are partially zippered in *trans*, Sec17 and Sec18 do not need HOPS to support the completion of fusion. The completion of SNARE zippering is promoted by Sec17 (Song et al., 2021), and Sec17 displaces HOPS from SNARE complexes (Collins et al., 2005; Schwartz et al., 2017). The interactions among the SNAREs, Sec17/αSNAP and Sec18/NSF are seen at a molecular level in the 20s complex, consisting of a 4-SNARE coiled coil anchored to membranes at their C-termini, surrounded by up to 4 Sec17/αSNAP molecules , and all capped at the membrane-distal end by Sec18/NSF (Zhao, 2015). A “*trans*-20s” (Song et al., 2021; Rizo et al., 2021) may generally drive fusion, since its elements are common to all exocytic and endocytic trafficking while the tethering complexes of other organelles are varied and often do not have the organelle’s SM protein as a tightly-bound subunit like Vps33 is in HOPS (Baker and Hughson, 2016). These findings support the model (Song et al., 2021) that HOPS only acts for tethering and to catalyze the initial phase of zippering in a Sec17-resistant manner. HOPS is specific for fusion at the vacuole/lysosome, but Sec17/αSNAP, Sec18/NSF, and SNAREs are general elements of exocytic and endocytic vesicular trafficking. Our current findings suggest that Sec17 and Sec18 support of zippering-bypass fusion may not be restricted to the vacuole/lysosome, but may broadly contribute to many SNARE-mediated fusion events.

## Methods

Reagents were purchased, and proteins purified, as described in Song et al., 2021.

### GST-PX constructions

DNA encoding the PX domain from the Qc SNARE Vam7 (Amino acyl residues 2-123) was amplified by PCR with CloneAMP HiFi PCR premix (Takara Bio USA, Mountain View, CA, USA). The amplified DNA fragment was cloned into BamHI and SalI digested pGST parallel1 vector (Sheffield et al.,1999) with an NEBuilder HiFi DNA Assembly kit (NEB, Ipswich, MA, USA).

For GST-PX

F AGGGCGCCATGGATCCGGCAGCTAATTCTGTAGGGAA

R AGTTGAGCTCGTCGACTATGGCTTTGACAACTGCAGGA

GST-PX was prepared as described (Fratti and Wickner, 2007).

Proteoliposome preparation and fusion assays were as described in Song et al. (2021). In brief, proteoliposomes were separately preincubated for 10 min at 27°C with EDTA and GTP, followed by addition of MgCl_2_, to load the Ypt7 with GTP. After separate preincubation for 10min at 27°C of both proteoliposome preparations and of mixtures of all soluble proteins (HOPS, GST-PX, Sec17, and Sec18/ATPγS) empty assay wells received in rapid succession 5μl of Ypt7/R proteoliposomes, 5μl of Ypt7/3Q proteoliposomes, and an 8μl mixture of all soluble components. FRET representing fusion was recorded each minute for 30 min, as described (Song et al., 2021), then the multiwell plate was withdrawn and 2μl of buffer, Sec17, Sec18, or a mixture of Sec17, Sec18, and Mg:ATPγS was added and the plate returned to the machine in time for the 31 minute time-point and those thereafter.

## Acknowledgements

We thank Amy Orr for expert technical assistance. This work was supported by NIGMS grant R35GM118037.

